# Hydrogel viscoelasticity modulates cell nascent extracellular matrix deposition

**DOI:** 10.1101/2025.06.06.658326

**Authors:** Matthew L. Tan, Eleanor M. Plaster, Avinava Roy, Haguy Wolfenson, Adam Abraham, Claudia Loebel

**Affiliations:** Department of Materials Science and Engineering, University of Michigan, Ann Arbor, MI 48109, USA; Department of Biomedical Engineering, University of Michigan, Ann Arbor, MI 48109, USA; Department of Genetic and Developmental Biology, Rappaport Faculty of Medicine, Technion, Haifa 352433, Israel; Department of Orthopedics, University of Michigan, Ann Arbor, MI 48109 USA; Department of Bioengineering,m University of Pennsylvania, Philadelphia, PA 19104, USA; Center for Precision Engineering for Health, University of Pennsylvania, Philadelphia, PA 19104, USA

**Keywords:** hydrogels, viscoelasticity, extracellular matrix deposition, mechanotransduction Abstract

## Abstract

Polymeric hydrogels are valuable platforms for determining how specific mechanical properties of native tissue extracellular matrix (ECM) regulate cell function. Recent research has focused on incorporating viscous and elastic properties into hydrogels to investigate cellular responses to time-dependent mechanical properties of the ECM. However, a critical aspect often overlooked is that cells continuously remodel their microenvironment in hydrogels, such as by the deposition of newly secreted (nascent) ECM. While this nascent ECM has been demonstrated to play a vital role in transmitting mechanical signals across various biological contexts, the mechanisms by which it regulates cellular function in response to time-dependent mechanical properties remain poorly understood. In this study, we developed an interpenetrating polymer network that enables independent control of viscous and elastic hydrogel properties. We show that cells cultured on high-viscosity hydrogels deposit increased nascent ECM which also correlates with enhanced hydrogel remodeling. Interestingly, higher nascent ECM deposition on high-viscosity hydrogels was decoupled from intracellular contractility. These results establish a relationship between hydrogel viscosity and nascent ECM deposition that may extend to diverse cell types and offer new insights into cell-hydrogel interactions.

## 1. Introduction

Polymeric hydrogels offer an adaptable range of mechanical properties to recreate aspects of native tissue and investigate their role in physiological processes. The high water content and tunable network structure of synthetic hydrogels have played critical roles in approaches to mimicking the native extracellular matrix (ECM) to study cell function *in vitro*.^[1,2]^ However, many of the initial studies were based on static and primarily elastic hydrogels^[3–5]^, which lack the dynamic and time-dependent nature of the ECM. To address this, recent strategies have focused on introducing viscosity and stress-relaxation into hydrogels, including through crosslinking with reversible bonds.^[6]^ Indeed, cells cultured atop or within these viscoelastic hydrogels actively remodel and respond to these time-dependent mechanical properties, which contributes to cell function.^[7–9]^ Critical to this process is the sensing and transmission of mechanical signals into cellular responses, known as mechanotransduction.^[10,11]^ Cells sense their extracellular environment through focal adhesions that connect to the nucleus via the actin cytoskeleton, facilitating the conversion of mechanical signals into intracellular responses.^[10–12]^ For example an increase in the elastic modulus of a hydrogel typically leads to an increase in intracellular contractility and consequent cell spreading.^[13]^ In addition, cells continuously remodel their surrounding ECM, including via the deposition of new ECM components such as proteins and sugars.^[14]^ This “dynamic reciprocity” between cells and their ECM has been shown to be critical in tissue homeostasis,^[15,16]^ yet the contributions of new ECM to hydrogel remodeling remains underexplored. In our previous work, we developed a metabolic labeling approach that enables visualization and identification of newly secreted (nascent) ECM (nECM).^[17,18]^ In fact, nECM deposition and remodeling by cells within 3D hydrogels have been shown to be critical for mechanotransduction in several cell types.^[17,19–23]^ Despite these advances in 3D hydrogels, how nECM deposition directs cellular function in response to time-dependent mechanical properties is largely unknown.

Here, we developed a phototunable interpenetrating polymer network (IPN) hydrogel to independently control viscous and elastic properties and applied this system to study the role of nECM deposition in cell function. To accomplish this, we used metabolic labeling and markers of intracellular contractility and focal adhesions to show the relationship between hydrogel viscosity, nECM deposition and cell mechanotransduction.

## 2. Results and Discussion

2.1. IPN hydrogels, which enable the independent tuning of viscous and elastic moduli. IPN hydrogels, were formed through simultaneous network formation of methacrylated-hyaluronic acid (MeHA) via kinetic chain-growth and norbornene-modified hyaluronic acid (NorHA) via thiol-ene step-growth of thiolated -guest (adamantane, Ad-SH) and -host (β-cyclodextrin, CD-SH) crosslinkers using visible light (**Figure 1A**).^[24]^ It is important to note that norbornenes react much faster with the thiols on CD and Ad than with methacrylates.^[25]^ Thus, the two networks are expected to be crosslinked primarily independent of each other.

**Figure 1.**
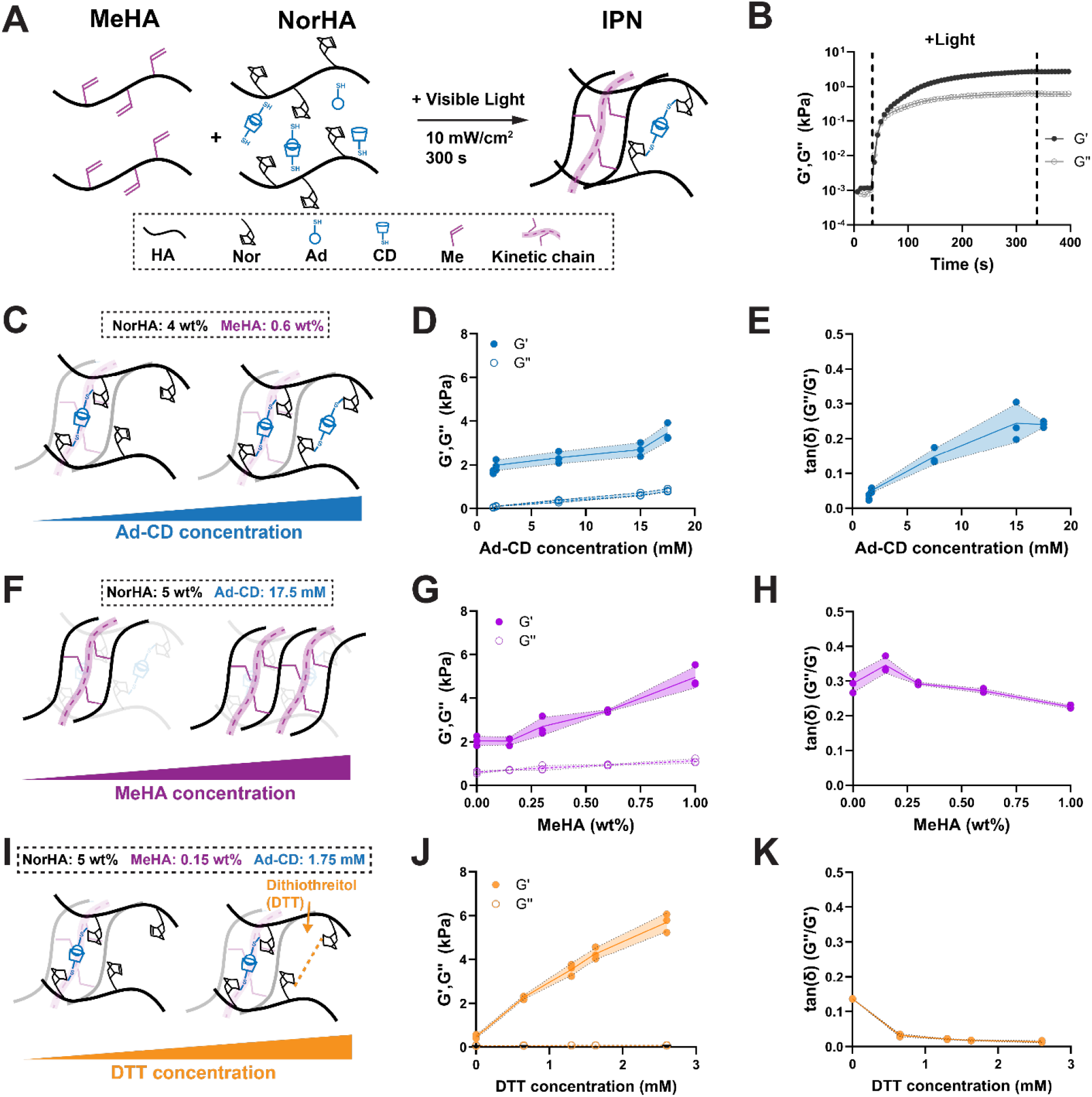
Interpenetrating polymer network (IPN) hydrogels enable independent tuning of viscous and elastic properties. **A**. Schematic illustrating the fabrication of IPN hydrogels based on methacrylated hyaluronic acid (MeHA), norbonene-modified hyaluronic acid (NorHA), and guest host crosslinkers (Ad-CD). **B**. Representative oscillatory time-sweep of IPN hydrogel photopolymerization at 15 mM Ad-CD, 4 wt% NorHA, and 0.15 wt% MeHA (storage modulus (G’), Loss modulus (G’’) at 10 rad/s and 0.1% strain). Area in between dotted lines indicate visible light exposure. **C**. Schematic illustrating the effect of increasing Ad-CD concentrations on IPN structure (constant NorHA: 4 wt%, MeHA: 0.6 wt%). **D**. Quantification of G’ and G’’ of IPN hydrogels as a function of Ad-CD concentration. **E**. Quantification of G’’/G’ (tanδ) as a function of Ad-CD concentration. **F**. Schematic illustrating the effect of MeHA concentration on IPN structure (constant NorHA: 5 wt%, Ad-CD: 17.5 mM). **G**. Quantification of G’ and G’’ as a function of MeHA concentration. **H**. Quantification of G’’/G’ (tanδ) as a function of MeHA concentration. **I**. Schematic illustrating the effect of increasing MeHA concentration on IPN structure (constant NorHA: 5 wt%, MeHA: 0.15 wt%, Ad-CD: 1.75 mM). **J**. Quantification of G’ and G’’ as a function of DTT concentration. **K**. Quantification of G’’/G’ (tanδ) as a function of DTT concentration. N = 3 hydrogels per group.

Oscillatory shear rheology showed a rapid increase in shear modulus (G’) and loss modulus (G’’) upon exposure to light with a gelation point at approximately 30 seconds that plateaued within 300 sec, indicating stable network formation (**Figure 1B**). To investigate the contributions of each component of the independent networks, we first altered the Ad-CD concentration while maintaining both the NorHA (4 wt%) and the MeHA (0.6 wt%) concentrations (**Figure 1C**). Increasing Ad-CD concentrations resulted in an approximately 15-fold increase in G’’ from 0.049 ± 0.013 kPa (1.5 mM Ad-CD) up to 0.834 ± 0.068 kPa (17.5 mM Ad-CD). In contrast, G’ increased by 2-fold from 1.672 ± 0.075 kPa (1.5 mM Ad-CD) to 3.472 ± 0.039 kPa (17.5 mM Ad-CD, **Figure 1D**). Calculating tanδ (ratio of G’’ to G’) demonstrated a near linear relationship between the molar concentration of Ad-CD and tanδ (**Figure 1E**). Notably, tanδ plateaued at 17.5 mM Ad-CD, most likely due to saturation of available norbornenes. Thus, increasing amounts of Ad-CD bonds within the network result in a more viscous hydrogel behavior that may be further modulated by increasing the amount of available norbornenes. Next, we modulated the MeHA concentration while keeping both the NorHA (5 wt%) and Ad-CD (17.5 mM) concentrations constant (**Figure 1F**). Note, that 5 wt% NorHA was chosen to provide an additional number of norbornenes for subsequent studies. Increasing MeHA concentrations increased G’ from 2.046 ± 0.218 kPa (0.0 wt%) to 4.958 ± 0.050 kPa (1.00 wt%) with minimal changes in G’’ (**Figure 1G**). As a result, changing the MeHA concentration induced little change in tanδ (between 0.293 and 0.226) (**Figure 1H**), especially when compared to changing Ad-CD concentrations (tanδ between 0.029 and 0.241). This data confirms the ability of MeHA to modulate IPN elasticity. The overall elasticity may further be changed independently of polymer concentration through the addition of dithiol crosslinkers, such as dithiothreitol (DTT), which result in additional covalent bonds between norbornene groups (**Figure 1I**).^[6]^ Thus, we next modulated the DTT concentration while maintaining the NorHA (5 wt%) and Ad-CD constant at a low concentration (1.75 mM) to provide free norbornene groups for DTT crosslinking. Increasing DTT increased G’ from 0.474 ± 0.085 kPa (0 mM DTT) to 5.701 ± 0.043 kPa (2.6 mM DTT) with minimal change in loss moduli across concentrations (**Figure 1J**). This resulted in a relatively small linear decrease in tanδ with increasing addition of DTT (**Figure 1K**). Taken together, these data demonstrate the ability to independently tune the viscous and elastic properties of IPN hydrogels by varying Ad-CD, MeHA and NorHA concentrations.

### 2.2. Hydrogel viscosity regulates nascent matrix deposition

Having shown tunable hydrogel viscoelasticity, we next selected low tanδ (0.032) hydrogels (i.e., low viscosity) and high tanδ (0.346) hydrogels (i.e., high viscosity) to investigate its role in regulating human mesenchymal stromal cell (hMSC) function and nascent matrix (nECM) deposition (**Figure 2A**). To enable cell adhesion, the hydrogel backbone was additionally modified with cell adhesive domains using thiolated RGD (2 mM) that binds to the norbornenes via thiol-ene reaction upon light exposure. First, we fabricated low tanδ (5 wt% NorHA, 1.00 wt% MeHA, 1.75 mM Ad-CD, 1.3 mM DTT) and high tanδ (5 wt% NorHA, 1.00 wt% MeHA, 17.5 mM Ad-CD, 0.0 mM DTT) hydrogels with the same G’ of 5 kPa. To assess nECM deposition, we used a previously developed metabolic labeling technique based on the incorporation of azide-modified methionine analogs (azidohomoalanine) into proteins as they are synthesized^[17,20]^. Upon culture, azide-modified proteins are then fluorescently labelled using cyto-compatible cycloaddition (DBCO-488) prior to fixation which prevents staining of intracellular nascent proteins. In addition, we co-stained for fibronectin which is known to be a critical component of the ECM.^[26,27]^ Upon 72 hours of culture, hMSCs showed elongated morphologies on both low tanδ and high tanδ hydrogels, surrounded by nECM that had similar structure to the fibronectin staining (**Figure 2B**). When comparing projected cell area, we observed a significant decrease for hMSCs cultured on high tanδ hydrogels compared to low tanδ hydrogels (**Figure 2C**). This observation aligns with previous reports using RGD modified hydrogels at similar elasticity,^[28,29]^ while others have demonstrated increased cell spreading for highly viscous hydrogels at lower G’.^[7,28,30,31]^ Quantification of projected nECM and fibronectin area showed no significant difference between low and high tanδ 5 kPa G’ hydrogels (**Figure 2D**). Notably, the projected area of fibronectin staining per cell area is much higher than the projected nECM area, which may be due to the adsorption of the fibronectin in the culture medium. These findings indicate that changes in cell area have little effect on nECM deposition on 5 kPa hydrogels. Based on previous studies using lower G’ hydrogels to probe mechanisms of viscosity-induced cell function,^[7,32]^ we next cultured hMSCs on softer 2 kPa G’ hydrogels. Similarly to 5 kPa, hMSCs showed elongated morphologies and deposited nECM and fibronectin on both low and high tanδ hydrogels (**Figure 2E**). Quantification of projected cell area also showed a decrease on high tanδ compared to low tanδ hydrogels (**Figure 2F**). Interestingly, nECM and fibronectin staining were significantly increased for hMSCs cultured atop high tanδ compared to low tanδ hydrogels (**Figure 2G**). These data show that high viscosity of relatively soft hydrogels promotes the deposition of nECM. Notably, our findings further suggest that nECM deposition does not directly rely on an increase in cell spread area.

**Figure 2.**
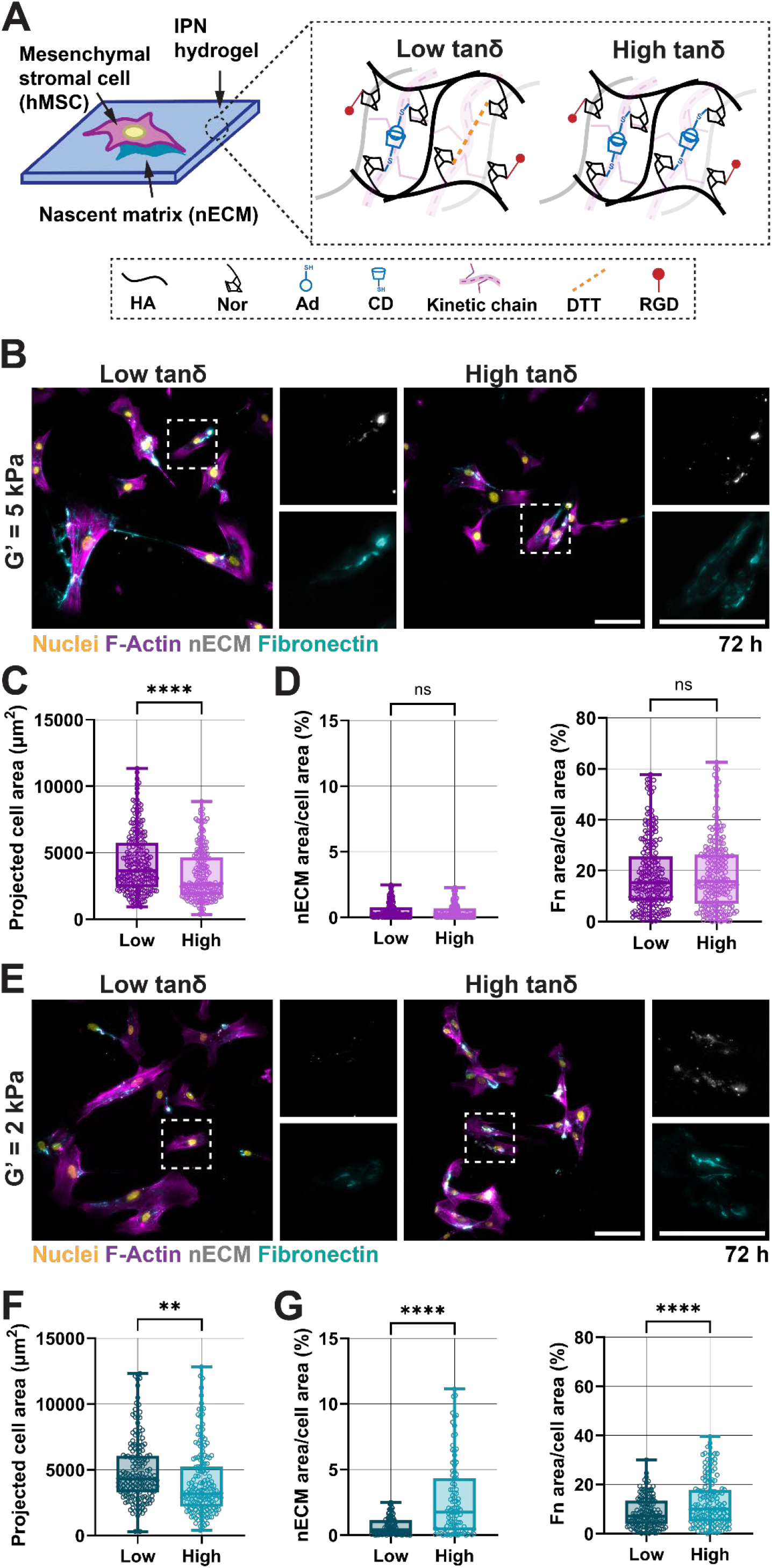
High tanδ soft hydrogels promote nECM deposition. **A**. Schematic illustrating the culture of hMSC atop low (tanδ = 0.032, 5 wt% NorHA, 1.75 mM Ad-CD, 4 mM RGD, 1.3 mM DTT) and high tanδ (tanδ = 0.346, 5 wt% NorHA, 17.5 mM Ad-CD, 4 mM RGD) hydrogels. **B**. Representative fluorescent images of hMSCs cultured for 72 hours (h) atop low and high tanδ hydrogels (G’ = 5 kPa, 1.00 wt% MeHA) Scale bars = 100 µm, 50 µm for insets. **C**. Quantification of projected hMSC spread area atop low and high tanδ hydrogels (G’ = 5 kPa, 72 h). N = 225, 213 cells total for low and high tanδ hydrogels respectively from 3 independent hydrogels **D**. Quantification of nECM and fibronectin staining of hMSCs atop low and high tanδ hydrogels (G’ = 5 kPa, 72 h). N = 225, 213 cells total for low and high tanδ hydrogels respectively from 3 independent hydrogels **E**. Representative fluorescent images of hMSCs cultured for 72 h atop low and high tanδ hydrogels (G’ = 2 kPa, 0.15 wt% MeHA). Scale bars = 100 µm, 50 µm for insets. **F**. Quantification of projected hMSC spread area atop low and high tanδ hydrogels (G’ = 2 kPa, 72 h). N = 169, 178 cells total for low and high tanδ hydrogels respectively from 3 independent hydrogels **G**. Quantification of nECM and fibronectin staining of hMSCs atop low and high tanδ hydrogels (G’ = 2 kPa, 72 h) N = 169, 178 cells total for low and high tanδ hydrogels respectively from 3 independent hydrogels. **p<0.01, ***p<0.001, ***p<0.0001, two-tailed student’s t-test with Welch’s correction.

### 2.3 High viscosity IPNs enable hydrogel remodeling via focal adhesions

Given that cells spread on IPN hydrogels by directly interacting with the polymer backbone (via the RGD motif), we next sought to determine the mechanism underlying cell spreading and nECM deposition on low and high tanδ (2 kPa G’) hydrogels. IPNs were fabricated with embedded fluorescent beads to map hydrogel deformation resulting from forces generated by hMSCs adhering and pulling on the polymer backbone. After 3 h of culture, hMSCs on both low and high tanδ induced some bead displacement at cell protrusions; however, no nECM was detected, presumably due to the relatively short culture time (**Figure 3A**). In contrast, after 72 h of culture, bead displacement increased across the entire cell area for hMSCs cultured atop high tanδ hydrogels surrounded by nECM (**Figure 3B**). Quantification of bead displacement confirmed little difference between low and high tanδ hydrogels after 3 h, whereas average displacement was significantly increased after 24 h and 72 h for hMSCs cultured on low tanδ compared to high tanδ hydrogels (**Figure 3C**). Thus, hMSCs remodel the IPN hydrogels with an increase in deformation for higher viscosity hydrogels, likely due to the reversible nature of Ad-CD bonds that permit increased polymer mobility.^[33]^ Interestingly, there was little overlap of deposited nECM and areas of high bead displacement.

**Figure 3.**
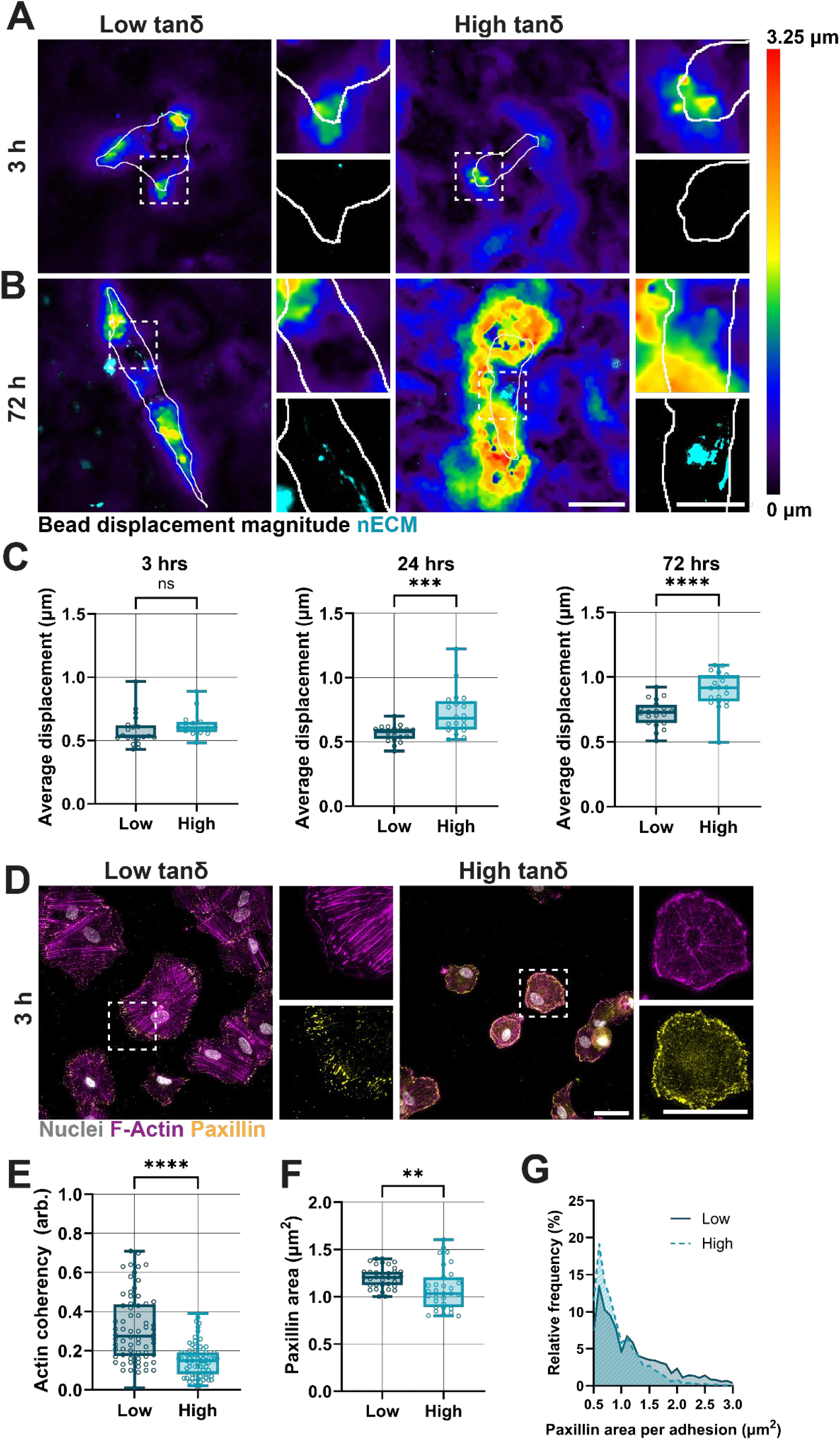
High viscosity IPNs increase hydrogel remodeling via focal adhesions. **A**. Representative images of bead displacement and nECM of hMSCs cultured for 3 h atop low and high tanδ hydrogels. Gray outline represents cell boundaries. Scale bars = 50 μm, 25 μm for insets. **B**. Representative images of bead displacement and nECM of hMSCs cultured for 72 h atop low and high tanδ hydrogels. Gray outline represents cell boundaries. Scale bars = 50 μm, 25 μm for insets. **C**. Quantification of average bead displacement by hMSCs cultured at low and high tanδ hydrogels for 3 h, 24 h, and 72 h. N = 20 cells total from 2 independent hydrogels **D**. Representative fluorescent images of F-Actin and paxillin of hMSCs cultured for 3 h at low and high tanδ hydrogels. Scale bars = 50 µm. **E**. Quantification of actin coherency of hMSCs cultured for 3 h at low and high tanδ hydrogels. N = 68, 71 cells total for low and high tanδ hydrogels respectively, from 3 independent hydrogels. **F**. Quantification of projected paxillin area of hMSCs cultured for 3 h atop low and high tanδ hydrogels. N = 35, 31 cells total for low and high tanδ hydrogels respectively, from 3 independent hydrogels. **G**. Quantification of relative frequency of single focal adhesion size (i.e., paxillin area) per hMSC cultured for 3 h atop low and high tanδ hydrogels. N = 35, 31 cells total for low and high tanδ hydrogels respectively, from 3 independent hydrogels. **p<0.01, ***p<0.001, ***p<0.0001, two-tailed student’s t-test with Welch’s correction.

In fact, orthogonal projection of nECM and fluorescent beads showed that nECM is deposited between the cell boundary and the hydrogel (**Figure S1**). This suggests that nECM is shielding the cells from the hydrogel and thus preventing hydrogel remodeling.

The ability of hMSCs to remodel the underlying hydrogel requires the expression of integrins and focal adhesions (paxillin) that enable mechanosensing and thus the remodeling of the actin cytoskeleton (F-Actin).^[34,35]^ After 3 h of culture atop low tanδ hydrogels, staining for F-Actin and paxillin showed strong actin fiber formation with aligned focal adhesions at the peripheral cell boundary whereas both actin fibers and focal adhesion showed more diffuse staining for hMSCs cultured atop high tanδ hydrogels (**Figure 3D**). F-Actin fiber orientation is often quantified by actin coherency^[36]^ which was significantly decreased for hMSCs cultured atop high tanδ hydrogels (**Figure 3E**), indicating lower intracellular contractility. Quantification of the average projected paxillin area also decreased for hMSCs cultured atop high tanδ hydrogels (**Figure 3F**). In addition, the higher relative frequency of small focal adhesions (i.e., <1.0 µm^2^) on high tanδ hydrogels (**Figure 3G**) further suggests potentially lower contractility and force generation by hMSCs atop viscous hydrogels.^[27,35,37]^ It is important to note that we also observed a significant decrease in the number of hMSCs that initially adhered to high tanδ hydrogels when compared to low tanδ hydrogels (**Figure S2**), which may explain the smaller focal adhesion size and actin coherence after 3 h of culture. Indeed, after 72 h, the average projected paxillin area was significantly higher for hMSCs cultured on high tanδ compared to low tanδ (**Figure S3**). Interestingly, we did not observe significant differences in actin coherence between low and high tanδ hydrogels after 72 h, which suggests that remodeling of high tanδ hydrogels does not rely on increased intracellular contractility. Taken together, although hMSCs are less likely to adhere to high tanδ hydrogels, longer culture periods enable hydrogel remodeling and the formation of more mature focal adhesions with little change in intracellular contractility between cells cultured on low and high tanδ hydrogels.

### 2.4 Intracellular contractility directs nECM deposition on high tanδ hydrogels

After having shown that hMSCs are smaller and initially (at the end of 3h) less contractile on high tanδ hydrogels, we next assessed the relationship between intracellular contractility and nECM deposition on these hydrogels. To increase intracellular contractility, we used the RhoA activator CN03, which has been shown to activate the formation of aligned actin stress fibers.^[38–41]^ hMSCs treated with CN03 and cultured for 3 h on high tanδ hydrogels showed similar F-Actin and paxillin staining when compared to untreated (CTRL) hMSCs (**Figure 4A**). However, quantification of cell area and actin coherency showed a significant increase upon CN03 treatment (**Figure 4B**). Average projected paxillin area showed minimal difference between low and high tanδ hydrogels (**Figure 4C)**. Quantification of relative frequencies of focal adhesions confirmed that there was little difference between the two groups (**Figure S4)**. Interestingly, on high tanδ hydrogels, CN03 treatment induced a significant increase in initial hMSCs adherence, reaching similar levels to low tanδ hydrogels (**Figure S5**). This may indicate that increased intracellular contractility supports initial cellular adhesion on high viscosity substrates independently of focal adhesion size.^[42]^ When treated for 72 h, no differences in actin coherency and focal adhesion area were observed (**Figure S6**), suggesting that CN03-inducing intracellular contractility may be most efficient during initial cell-hydrogel interactions.

**Figure 4.**
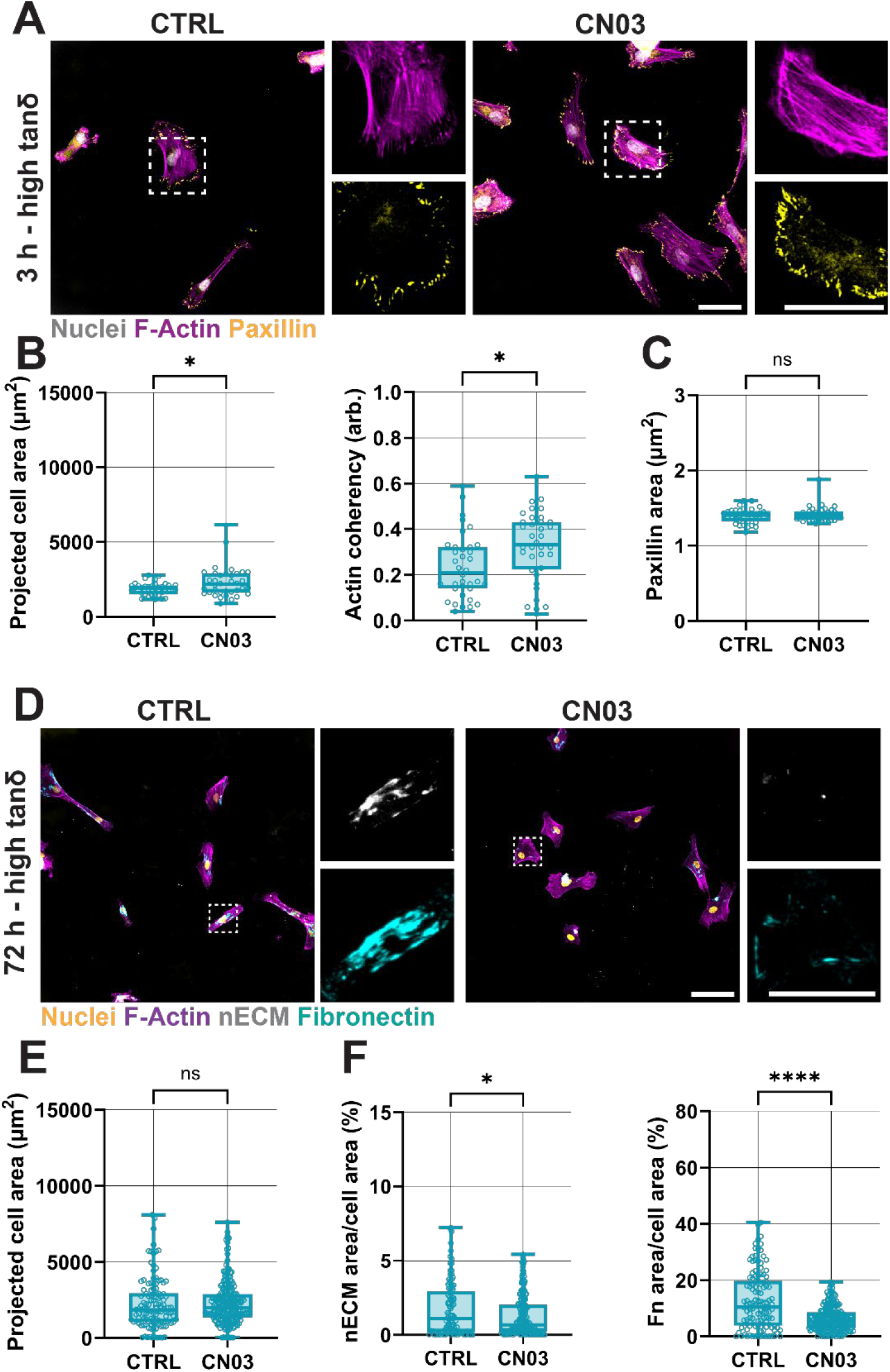
Intracellular contractility decreases nECM deposition on high tanδ hydrogels. **A**. Representative fluorescent images of F-Actin and paxillin of hMSCs cultured for 3 h atop high tanδ IPN hydrogels without (CTRL) or with 1 ug/mL CN03. Scale bars = 50 μm **B**. Quantification of projected cell area and projected actin coherency of hMSCs cultured for 3 h atop high tanδ IPN hydrogels without (CTRL) or with 1 ug/mL CN03. N = 36 cells total for both conditions from 3 independent hydrogels **C**. Quantification of projected paxillin area of hMSCs cultured for 3 h atop high tanδ IPN hydrogels without (CTRL) or with 1 ug/mL CN03 (see single focal adhesion frequency in Figure S4). N = 36 cells total for both conditions from 3 independent hydrogels. **D**. Representative fluorescent images of F-Actin, nECM and fibronectin of hMSCs cultured for 72 h atop high tanδ IPN hydrogels without (CTRL) or with 1 ug/mL CN03. Scale bars = 100 µm, 50 µm for insets. **E**. Quantification of projected cell area of hMSCs cultured for 72 h atop high tanδ IPN hydrogels without (CTRL) or with 1 ug/mL CN03. N = 120, 232 cells total for CTRL and CN03 respectively, from 3 independent hydrogels. **F**. Quantification of projected nECM and fibronectin staining of hMSCs cultured for 72 h atop high tanδ IPN hydrogels without (CTRL) or with 1 ug/mL CN03. N = 120, 232 cells total for CTRL and CN03 respectively, from 3 independent hydrogels. *p<0.05, ****p<0.0001, two-tailed student’s t-test with Welch’s correction.

Next, we assessed how the CN03-mediated increase in intracellular contractility directs nECM deposition on high tanδ hydrogels. After 72 h of culture, hMSCs remained elongated on high tanδ hydrogels in both CTRL and CN03-treated conditions but showed a strong reduction in both nECM and fibronectin staining in response to CN03 (**Figure 4D**).

Interestingly, quantification of projected cell area showed minimal differences between CTRL and CN03-treated hMSCs on high tanδ hydrogels (**Figure 4E**), corroborating the actin coherency and focal adhesion area measurements (Figure S6). In contrast, quantification of nECM and fibronectin showed significant decrease for CN03-treated hMSCs (**Figure 4F**).

These findings further confirm that cell area itself is not a strong indicator of nECM deposition. Interestingly, RhoA-mediated intracellular contractility has often been associated with increased ECM deposition, including the assembly of fibronectin into fibers.^[27,43]^ However, other reports have demonstrated that increasing intracellular contractility may also result in decreased ECM assembly such as through increased ECM degradation.^[44,45]^ While the mechanisms governing intracellular contractility and nECM deposition remain to be investigated, our findings suggest that there is a relationship between viscosity, intracellular contractility and nECM deposition.

## 3. Conclusion

In this work we engineered a tunable polymeric hydrogel network to independently tune the viscous and elastic properties and its contributions to the deposition of newly secreted proteins. Viscoelastic hydrogels have been instrumental in investigating how ECM mechanical properties regulate cell function. Often, previous work has focused on direct interactions of cells with the underlying hydrogel,^[3,7]^ which underestimates the potential importance of newly deposited ECM proteins. Indeed, we and others have shown that cells, upon embedding into 3D hydrogels, quickly deposit nECM which contributes to mechanosensing of viscoelasticity.^[17,19]^ Yet, a direct relationship between viscoelasticity and nECM deposition atop 2D hydrogels has not been established. By using metabolic labeling and fluorescent tracking of the underlying IPN hydrogels, we showed that high viscosity hydrogels enhance nECM deposition which is associated with hydrogel remodeling. Our work further suggests that increasing intracellular contractility may not necessarily be related to higher nECM deposition atop viscous hydrogels. We anticipate that further modulating pathways of intracellular contractility such as YAP/TAZ nuclear translocation^[46]^ and focal adhesion kinase^[47,48]^ may be used to study the role of viscosity and mechanosensing in nECM deposition.

While this study focused on RGD as cell-adhesive peptide, other binding motifs such as collagen type 1-derived (GFOGER) or laminin-derived (IKVAV) may be used to further study the cell function and nECM deposition as a function of viscous properties.^[28]^ Finally, although we have shown that hMSCs on high viscosity hydrogels deposit increased nECM, the spatial distribution is distinct from cells within 3D hydrogels. While these difference may in part be attributed to the effect of cell polarity and nECM release into the media for 2D culture, further studies are required to probe whether nECM alters mechanosensing of the underlying hydrogel^[14,17,19]^ Taken together, the IPN hydrogel and metabolic labeling technique described herein provides means as a tunable hydrogel system for studying the role of viscous and elastic properties in directing cell function and nECM deposition, which is extendable to other cell types.

## 4. Experimental Section

### Cell culture

Bone marrow-derived human mesenchymal stromal cells (hMSCs) were isolated from fresh bone marrow as previously described.^[49]^ Briefly, fresh bone marrow (Lonza) was strained through a 70 µm cell strainer and diluted 1:4 with phosphate buffered saline (PBS) and layered on top of Histopaque-1077 (Sigma) for density gradient centrifugation. Layered bone marrow was then centrifuged at 800 RCF for 20 minutes, and the mononuclear cell layer was collected and rinsed with α-MEM (Invitrogen) + 10% fetal bovine serum (FBS). Mononuclear cells were seeded on tissue culture plastic flasks in α-MEM + 10% FBS, 1% penicillin/streptomycin (P/S), 5 ng ml^-1^ basic fibroblast growth factor at 37°C, 5% CO2 until colonies were approximately 80% confluent. hMSCs were subsequently collected and stored in liquid nitrogen until use. For routine cell culture, hMSCs were maintained in α-MEM + 10% FBS, 1% P/S and harvested at 80% confluency. For hydrogel culture, hMSCs were maintained in glutamine, cystine and methionine free DMEM (Invitrogen) supplemented with 10% FBS, 1% P/S, 50 µM azidohomoalanine (AHA) (Vector Labs), 201 µM cystine, 2 mM GlutaMAX, 110 mg mL^-1^ sodium pyruvate, 50 µM methionine, and 50 µg mL^-1^ ascorbate 2-phosphate (AHA media). hMSCs were used between passages 1 – 3.

### Polymer synthesis and characterization

NorHA was synthesized as previously described.^[50]^ Briefly, sodium hyaluronate (HA) was solubilized in 2-(N-morpholino)ethanesulfonic acid buffer (pH 5.5) at 1% w/v. 4-(4,6-Dimethoxy-1,3,5-triazin-2-yl)-4-methylmorpholinium chloride (DMTMM) was added to solubilized HA, followed by dropwise addition of 5-norbornene-2-methylamine (Nor). The reaction was carried out for 25 h at 30°C to obtain an approximate degree of substitution of 36%. To isolate polymer, saturated sodium chloride was added, followed by precipitation with 200 proof ethanol. Polymer was then collected via vacuum filtration, washed, resolublized in deionized water, and dialyzed for 3 days through 6-8 kDa tubing before lyophilization and storage at -20°C. MeHA was synthesized as previously described.^[51]^ Briefly, HA was dissolved in deionized water, and the pH was adjusted to 8.5 with 1 M sodium hydroxide. Methacrylic anhydride (MA) was added dropwise while stirring and maintaining a pH of 8.5 – 9.5 and ice-cold conditions throughout the 8 h reaction. Polymer was dialyzed against deionized water for 5 days, before lyophilization and storage at - 20°C. Degree of substitutions for NorHA and MeHA were determined through H-NMR as previously described^[50,51]^ as approximately 36% and 30% respectively.

### Hydrogel fabrication

IPN hydrogels were fabricated as previously described.^[24]^ Briefly, MeHA and NorHA were dissolved in PBS using a water-bath sonicator. Stock solutions of 1-adamantanethiol (Ad-SH) (Sigma) and 6-mercapto-6-deoxy-β-cyclodextrin (CD-SH) (Crysdot) were prepared by dissolving compounds in DMSO at a concentration of 400 mM. To prepare a stock concentration of guest-host crosslinker, Ad-SH and CD-SH were combined in a 1:1 ratio (200 mM) and diluted with DMSO to 175 mM. Hydrogel precursor solutions were obtained by combining MeHA (0.15 – 1.00 wt %), NorHA (5 wt%), guest-host crosslinker (1.50 – 17.5 mM), thiolated RGD (4 mM), DTT (0 – 1.3 mM), and the photoinitiator lithium phenyl-2,4,6-trimethylbenzoylphosphinate (0.05%, LAP, Colorado Photopolymer Solutions). Precursor solutions were mixed with a pipette tip, vortexed for 3 minutes, and briefly centrifuged to remove bubbles. Precursor solutions were pipetted as 15 µL droplets onto glass coverslips treated with Sigmacote (Sigma), and a thiolated 12 mm round coverslip was placed on top of each droplet. Precursor solutions were subsequently photopolymerized with visible light (455 nm) for 5 minutes at 10 mW cm^-2^. Polymerized hydrogels were carefully removed from the Sigmacote-treated coverglass and placed into 24-well plates before rinsing 3 times with PBS. Hydrogels were then sterilized in a UV-sterlization chamber for 20 minutes before a final wash with PBS. Hydrogels were stored for at least 24 hours at 4°C before cell seeding.

### Shear rheology mechanical characterization

IPN hydrogel mechanical characterization was performed using an HR 30 Discovery Hybrid Rheometer (TA Instruments) with a 20 mm-diameter, 2° cone and plate geometry, and a 57 µm gap. IPN precursor solutions were pipetted onto a UV-attachment bottom geometry and exposed to visible light as described above to polymerize hydrogels during shear rheology. Elastic storage and viscous loss moduli were measured using through oscillatory time sweeps at 10 rad s^-1^, 0.1% strain.

### Hydrogel cell seeding and small molecule inhibition

To seed cells onto IPN hydrogels, hydrogels were first incubated in AHA media at 37°C for 30 minutes. hMSCs were harvested with 0.05% Trypsin-EDTA (Invitrogen) and resuspended in AHA media. Hydrogels were seeded at a cell density of 3000 cells cm^-2^ and cultured for 3, 24, and 72 h. Media was refreshed every other day.

To induce cell contractility, a stock solution of rho activator II (CN03) (Cytoskeleton, Inc.) was prepared by dissolving compound in sterile water to achieve a concentration of 200 µg mL^-1^. CN03 stock was diluted in AHA media to a final concentration of 1 µg mL^-1^ and added to a cell pellet for subsequent cell seeding. CN03 containing AHA media was refreshed daily.

### Nascent matrix labeling and immunofluorescence staining

Metabolic labeling and nascent matrix labeling were performed as previously described.^[18]^ Briefly, hydrogel cultures were rinsed with PBS containing calcium and magnesium, and incubated in 30 µM AZDye 488 dibenzocyclooctyne (DBCO) (Vector Labs) with 2% bovine serum albumin (BSA) in PBS containing calcium and magnesium at 37°C. Hydrogel cultures were rinsed 3 times with PBS before subsequent fixation in 4% paraformaldehyde (PFA) for 20 minutes. For focal adhesion staining, hydrogels were fixed/permeabilized with microtubule stabilizing fixation buffer (0.1 M 1,4-Piperazinediethanesulfonic acid, 1 mM ethylene glycol-bis(2-aminoethylether)-N,N,N’,N’-tetraacetic acid, 1 mM magnesium sulfate, 4% w/v polyethylene glycol, 1% v/v Triton X-100, 2% PFA). After fixation, hydrogels were rinsed 3 times with PBS and blocked in 2% BSA in PBS (blocking buffer) for 1 h. Primary antibodies were diluted in blocking buffer and added to hydrogel cultures for staining overnight at 4°C. Primary antibodies include anti-fibronectin (1:200, Sigma F6140), anti-laminin (1:200, Abcam ab11575), and anti-paxillin (1:200, BD Biosciences 610052). After washing 3 times with PBS, secondary antibodies and stains (AlexaFluor 647, Phalloidin 568 for actin staining, Hoechst 33342 for nuclear staining) were diluted in blocking buffer and added to hydrogel cultures for staining for 1 h at room temperature. Hydrogels were washed 3 times with PBS before imaging and/or storage at 4°C.

### Imaging and image analysis

Images of cells stained for nascent matrix and matrix proteins were acquired on a Leica DMi8 THUNDER widefield microscope with a 25x water immersion objective or a Cytation C10 (Agilent) with a 20x air objective, while images of cells stained for paxillin were acquired using a 40x water immersion objective or 40x air objective on the Leica and Cytation, respectively.

All image quantification was performed using ImageJ. For matrix and cell spread area quantification, 10 fields of views were obtained per hydrogel. Individual cells were segmented based on f-actin channel using Otsu’s thresholding, and regions of interests (ROIs) were created of each cell perimeter. Cell spread area was calculated based off ROI area, while nascent matrix and matrix protein positive staining area were determined in each ROI using a manually set threshold that was kept consistent between all experiments. Matrix staining positive area was subsequently normalized to cell spread area and each cell analyzed plotted as a single data point. For focal adhesion analysis, individual cells were cropped from 10 fields of views (approximately 10 cells per hydrogel). The paxillin channel was processed through background subtraction (rolling ball radius of 20 pixels), gaussian blur (sigma of 1 pixel), and a top hat filter (radius of 4 pixels) to isolate individual focal adhesions (paxillin) and reduce background signal. Individual focal adhesions were identified with an area greater than 0.75 µm^2^, and individual focal adhesion area was calculated and averaged per cell (each data point = 1 cell). Histograms of focal adhesion area were generated across all cells analyzed for each experiment. For actin organization analysis, the f-actin channel was processed using the OrientationJ plugin’s “Dominant Direction” function, and coherency was calculated per cell. All quantification was carried out using automated ImageJ scripts.

### Bead displacement imaging and quantification

To assess cell contractility, 0.2 μm nominal diameter red fluorescent microspheres (Thermo Fisher Scientific) sonicated in a water bath for 5 minutes and added to hydrogel precursor solutions at a 1:100 dilution. Thiolated 12 mm coverslips were affixed to the bottom of 24-well glass plates (Cellvis) with Norland Optical Adhesive 61 and UV cured at 365 nm for 1 minute. Well plates were placed in a UV-chamber for 20 minutes to finish curing. To form hydrogels, precursor solution was deposited on top of affixed thiolated coverslips and covered with a Sigmacote-treated 12 mm coverslip (30 μm thickness hydrogel). Hydrogels were photopolymerized as described above and sterilized in a UV-chamber for 20 minutes before washing with PBS overnight. Cells were seeded onto microsphere-containing hydrogels in AHA medium at 1,000 cells cm^-2^ for 3, 24, and 72 h. Nascent matrix labeling was performed as described above, and Fluorobrite DMEM (Invitrogen) + 10% FBS, 2 mM GlutaMax was added to hydrogels. Hydrogel containing wellplates were placed in a Cytation C10 imager at 37°C, 5% CO2. Phase contrast images of cells and fluorescent images of microspheres and nascent matrix was obtained using a 40x air objective. Images were obtained before and after the addition of sodium dodecyl sulfate (0.5% final concentration in PBS) to obtain contracted and relaxed position of fluorescent microspheres respectively. Bead displacement magnitude was calculated and plotted using a particle image velocimetry plugin in ImageJ.^[52]^

### Statistical analysis

All statistical analysis were performed using GraphPad Prism (version 10). Each experiment was performed at least three times independently using hMSCs isolated from one donor, unless specified otherwise. Comparisons between two groups were performed with an unpaired, two-tailed Student’s t-test with Welch’s correction. Comparisons between three or more groups were performed with a Brown-Forsythe one-way ANOVA, and multiple comparisons were performed with Dunnett’s T3 multiple comparison tests.

## Supporting Information

Supporting Information is available from the Wiley Online Library or from the author.

## Acknowledgements

This work was supported by funding from the NIH (R00-HL151670 to C.L., NHLBI T32 HL007749 to M.L.T), the David & Lucile Packard Foundation (to C.L) and the Michigan-Israel Partnership For Research And Education (to C.L. and H.W.).

