## Supplement for "Hydrogel viscoelasticity modulates cell nascent extracellular matrix deposition"

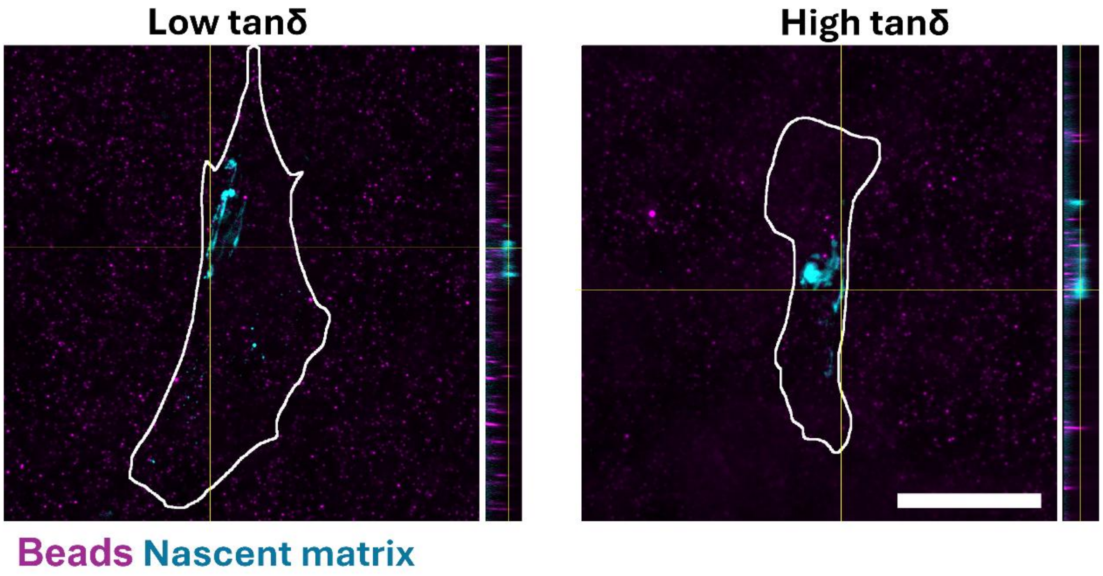


**Figure S1.** Representative confocal slice and orthogonal projection of cells (cell boundary marked) seeded on bead-laden low and high tanδ hydrogels after 72 h of culture. Scale bar = 50 µm

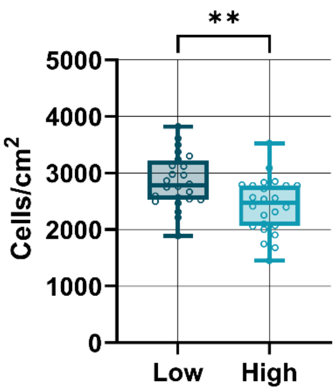


**Figure S2.** Number of cells per region of interest attached to low and high tanδ hydrogels 3 h post-seeding, wash, and fixation. N = 8 regions per hydrogel for 3 independent hydrogels, **$p<0.01$, two-tailed student's t-test with Welch's correction.

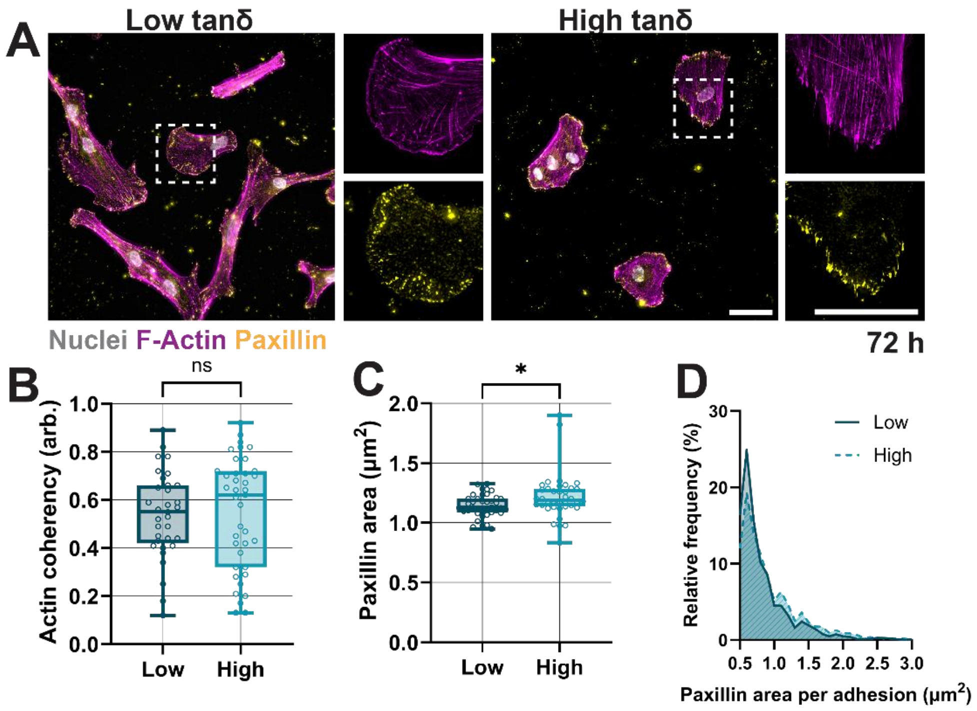


**Figure S3.** 72 h timepoint of focal adhesion and actin analysis. **A.** Representative fluorescent images of F-Actin and paxillin of hMSCs cultured for 72 h at low and high tanδ hydrogels. Scale bars = 50 µm. **B.** Quantification of actin coherency of hMSCs cultured for 72 h at low and high tanδ hydrogels. N = 33, 39 cells total for low and high tanδ hydrogels respectively, from 3 independent hydrogels. **C.** Quantification of projected paxillin area of hMSCs cultured for 72 h atop low and high tanδ hydrogels. N = 38 cells total from 3 independent hydrogels. **D.** Quantification of relative frequency of single focal adhesion (i.e., paxillin area) per hMSC cultured for 72 h atop low and high tanδ hydrogels. N = 38 cells total from 3 independent hydrogels. **p<0.01, ***p<0.001, ***p<0.0001, two-tailed student's t-test with Welch's correction.

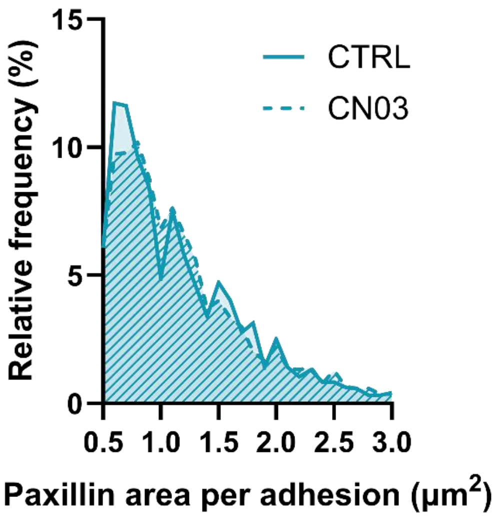


**Figure S4:** Quantification of relative frequency of single focal adhesion (i.e., paxillin area) per hMSC cultured for 3 h atop non-treated (CTRL) or CN03 treated (CN03) high tanδ hydrogels. N = 36 cells total from 3 independent hydrogels.

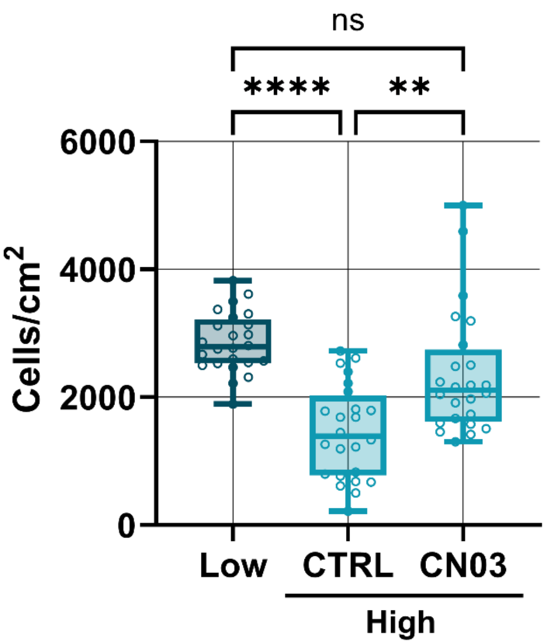


**Figure S5.** Number of cells per region of interest attached to low tan δ hydrogels without CN03 (Low) and high tanδ hydrogels without (CTRL) or with CN03 treatment (CN03) 3 h post-seeding, wash, and fixation. N = 8 regions per hydrogel for 3 independent hydrogels, ns = non-significant, **p<0.01, ***p<0.001, Brown-Forsythe and Welch ANOVA with Dunnett T3 test for multiple comparisons.

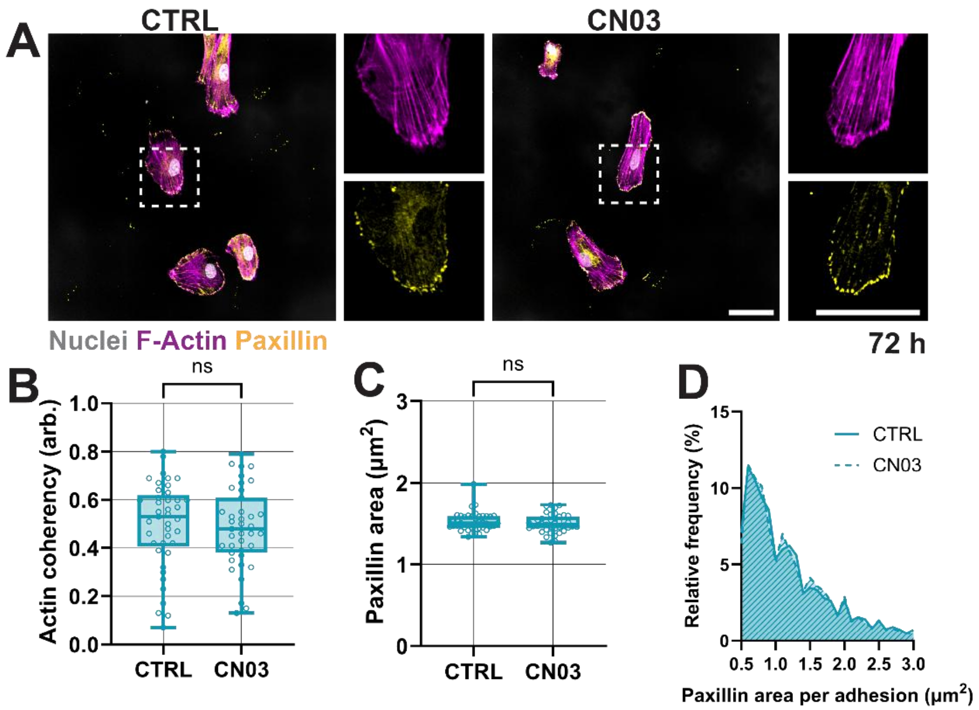


**Figure S6**. 72 h timepoint of focal adhesion and actin analysis with or without CN03 treatment. **A.** Representative fluorescent images of F-Actin and paxillin of hMSCs cultured for 72 h on high tanδ hydrogels without (CTRL) or with CN03 treatment (CN03). Scale bars = 50 µm. **B.** Quantification of actin coherency of hMSCs cultured for 72 h on high tanδ hydrogels without (CTRL) or with CN03 treatment (CN03). N = 41, 39 cells total for CTRL and CN03 respectively, from 3 independent hydrogels. **C.** Quantification of projected paxillin area of hMSCs cultured for 72 h on high tanδ hydrogels without (CTRL) or with CN03 treatment (CN03). N = 41, 39 cells total for CTRL and CN03 respectively, from 3 independent hydrogels. **D.** Quantification of relative frequency of single focal adhesion (i.e., paxillin area) per hMSC cultured for 72 h atop low and high tanδ hydrogels. N = 41, 39 cells total for CTRL and CN03 respectively, from 3 independent hydrogels. ns = non-significant, two-tailed student's t-test with Welch's correction.
